# Chromosome scale reference genome of Cluster bean (*Cyamopsis tetragonoloba* (L.) Taub.)

**DOI:** 10.1101/2020.05.16.098434

**Authors:** Kishor Gaikwad, G. Ramakrishna, Harsha Srivastava, Swati Saxena, Tanvi Kaila, Anshika Tyagi, Priya Sharma, Sandhya Sharma, R Sharma, HR Mahla, SV Amitha Mithra, Amol Solanke, Pritam Kalia, AR Rao, Anil Rai, TR Sharma, NK Singh

## Abstract

Clusterbean (*Cyamopsis tetragonoloba* (L.) Taub.), also known as Guar is a widely cultivated dryland legume of Western India and parts of Africa. Apart from being a vegetable crop, it is also an abundant source of a natural hetero-polysaccharide called guar gum or galactomannan which is widely used in cosmetics, pharmaceuticals, food processing, shale gas drilling etc. Here, for the first time we are reporting a chromosome-scale reference genome assembly of clusterbean, from a high galactomannan containing popular guar cultivar, RGC-936, by combining sequenced reads from Illumina, 10x Chromium and Oxford Nanopore technologies. The initial assembly of 1580 scaffolds with an N50 value of 7.12 Mbp was generated. Then, the final genome assembly was obtained by anchoring these scaffolds to a high density SNP map. Finally, a genome assembly of 550.31 Mbp was obtained in 7 pseudomolecules corresponding to 7 chromosomes with a very high N50 of 78.27 Mbp. We finally predicted 34,680 protein-coding genes in the guar genome. The high-quality chromosome-scale cluster bean genome assembly will facilitate understanding of the molecular basis of galactomannan biosynthesis and aid in genomics-assisted breeding of superior cultivars.

## Introduction

Clusterbean (*Cyamopsis tetragonoloba* (L.) Taub.), also known as guar^1^ is a member of Leguminosae family. The common name clusterbean is attributed to its pods which appear in clusters. Previous reports suggest that guar originated in Africa and later spread to the entire South Asian region. In India and Pakistan, clusterbean is cultivated since ancient times for its tender pods which are used as fresh vegetable and the remaining plant serves as fodder^2^. Clusterbean is a climate-resilient annual legume and a high potential alternative crop in the marginal lands of arid and semi-arid regions^3^. The genus *Cyamopsis* includes four species i.e., one cultivated *C. tetragonoloba* (L.) Taub., two wild relatives *C. serrata Schinz*, and *C. senegalensis Guill*&*Perr, and C. dentate Tarre*, and an interspecies hybrid of *C. serrata and C. senegalensis*^4^. A mature clusterbean seed is composed of three parts: germ (43-47%), endosperm (35-42%), and seed coat (14-17%). About 80-90% of the endosperm is composed of highly viscous water-soluble hetero-polysaccharide called gaur gum (or) galactomannan, having a 1:2 ratio of galactose to mannose^5^. Guar gum is extensively utilized as natural thickener, emulsifier and stabilizers in the food, textile, paper, petroleum and pharmaceutical industry with increasing global demand^6–9^. With the annual production of ∼1-1.25 million tons of clusterbean seeds, India accounts for 80% of the global production, with several other countries, like Pakistan, United States, China, Australia and Africa contributing the rest. About 45% of total world demand is due to industrial application of guar gum^10^. Apart from being a rich source of commercial product like gum, clusterbean is also a highly nutritious legume crop, predominantly composed of protein (18%) and dietary fiber (32%)

Earlier cytogenetic studies in clusterbean revealed that ∼580.9 Mb of the genome is organized in 2n=14 number of chromosomes^11,12^. Despite the considerable industrial importance, only a few studies have been carried out at genome level which includes genome size estimation (cultivated vs. wild type)^12^, plastid genome sequencing (Chloroplast)^13^, transcriptome analysis^14,15^, small RNA sequencing^16^ to identify novel miRNA associated with galactomannan biosynthesis as well as genetic diversity analysis based on SSRs^17^, mostly from our group. Therefore, it was necessary to sequence a chromosome-scale high-quality reference genome to understand the molecular basis of galactomannan biosynthesis, synteny with other legumes and discovery of genes for other important traits. This will also enhance cluster bean genetic improvement via genomics assisted breeding.

## Experimental Methods and Results

### Plant sample collection, genomic library preparation and sequencing

Seeds of the pure homozygous cluster bean variety, ‘RGC-936’, obtained from ICAR-CAZRI were sown in pots at ICAR-NIPB, New Delhi, India. Leaf samples were collected and snap frozen in liquid nitrogen and stored at -80°C till further use. The genomic DNA was extracted using CTAB method^18^ and the integrity and quantity of DNA were tested by separating the DNA on a 0.8% agarose gel and DeNovix DS-11 spectrophotometer, respectively. The high-quality DNA was used for genome sequencing by Illumina and HMW DNA was used for 10X Genomics and Oxford Nanopore sequencing. Similarly leaf samples of a F2 population (RGC 936 x CAZRI-15-3-8) were processed for GBS sequencing (Illumina).

### Genome sequencing

In the present study, we selected both short and long-read sequencing methods such as Illumina, 10X Genomics and Oxford Nanopore sequencing technology (ONT) for clusterbean genome sequencing.

For Illumina short-read sequencing, high-quality genomic DNA was randomly fragmented by the M220 Focused-ultra sonicator system (Covaris Inc, USA). Two genomic DNA libraries of 500-1000bp insert size were prepared using the TruSeq DNA PCR-Free Sample Preparation Kit, as per the manufacturer’s guidelines. To realize sequence variation and high genome coverage (length), two separate Mate-Pair (MP) libraries of 3Kb and 7Kb insert size were prepared using Nextera Mate-Pair Sample Preparation Kit (Illumina, San Diego, CA). Both the Illumina PE and MP libraries were sequenced on Illumina HiSeqX-ten platform which produced 16.09Gbp and 42.3Gbp of (2×150bp) sequencing data respectively.

For 10X Genomics sequencing, a total of 8 Gemcode libraries were prepared from high-quality DNA fragments longer than 50Kb, using the Chromium instrument. Sequencing of these libraries was performed on Illumina, HiSeqX-ten platform, generating 2×150bp reads; resulting in a total of 92.767 Gbp of 10X Genomics linked read raw sequencing data.

For long-read Nanopore sequencing, genomic DNA was size-selected using BluePippin BLF7510 cassette (Sage Science) and high-pass mode (> 20 kb) and library was prepared using Oxford Nanopore Technologies (ONT) standard ligation sequencing kit SQK-LSK109 following the SQK-MAP005 PromethION protocol. Two libraries were prepared and sequenced using PromethION. A total of 50.64 Gbp of Nanopore sequencing data was generated.

As a result, we generated a total of 201.8 Gbp raw sequencing data corresponding to 366.73X genomic coverage (depth), of cluster bean genome (543.22 Mbp estimated genome size using *k*-mer frequency distribution analysis). Details of sequencing data are represented in **Table 1**.

**Table 1:** Raw sequencing data generated for cluster bean genome assembly.

| Library type | Reads length<br>(bp) | Insert size<br>(bp) | No. of<br>libraries | No. of<br>Reads | Sequence output<br>(bp) | Genome<br>coverage |
| --- | --- | --- | --- | --- | --- | --- |
| *Illumina pair-ended/mate-pair sequencing* |  |  |  |  |  |  |
| PCR-free | 2x150 | 470 | 1 | 54,304,615 | 10,427,940,295 | 18.95 X |
| PCR-free | 2x150 | 800 | 1 | 29,507,608 | 5,666,395,556 | 10.30 X |
| Mate-Pair | 2x160 | 3000 | 1 | 84,083,542 | 24,261,975,442 | 44.09 X |
| Mate-Pair | 2x160 | 7000 | 1 | 63,517,764 | 18,046,975,201 | 32.79 X |
| *10 x Genomics Linked Reads* |  |  |  |  |  |  |
| Chromium | 2x150 |  | 8 | 309,223,705 | 92,767,111,500 | 168.57 X |
| *Oxford Nanopore* |  |  |  |  |  |  |
| PromithION | 2 |  |  | 6,712,699 | 50,647,765,074 | 92.03 X |
| **Total** |  |  |  |  | **201,818,163,068** | **366.73 X** |

### Genome size estimation

In the present study, the cluster bean genome size was estimated using the *k-*mer frequency distribution analysis of short reads sequencing data with a *k*-mer size of 31. The 31-mer abundance was calculated using Jellyfish^19^ v2 and the cluster bean genome size was then estimated using Genomescope software^20^. Total 1,187,123,081 *k*-mers (31-mer) were counted and their frequency distributions were analyzed. In the 31-mer frequency distribution histogram, the main peak was observed at depth of 65 corresponding to homozygous haploid sequences. A small peak was also observed at half of the main peak depth representing heterozygous fraction of the genome (**Figure 1a**). The *k*-mer frequency distribution histogram produced by Jellyfish was then subjected to Genomescope (http://qb.cshl.edu/genomescope/genomescope2.0/) for estimating size and heterozygosity of the genome. Thus, cluster bean genome size was estimated to be 543.2 Mb (543,218,592 bp), and the fraction of heterozygosity in cluster bean genome estimated to be in the range of 0.70 - 0.71%.

**Figure 1:**
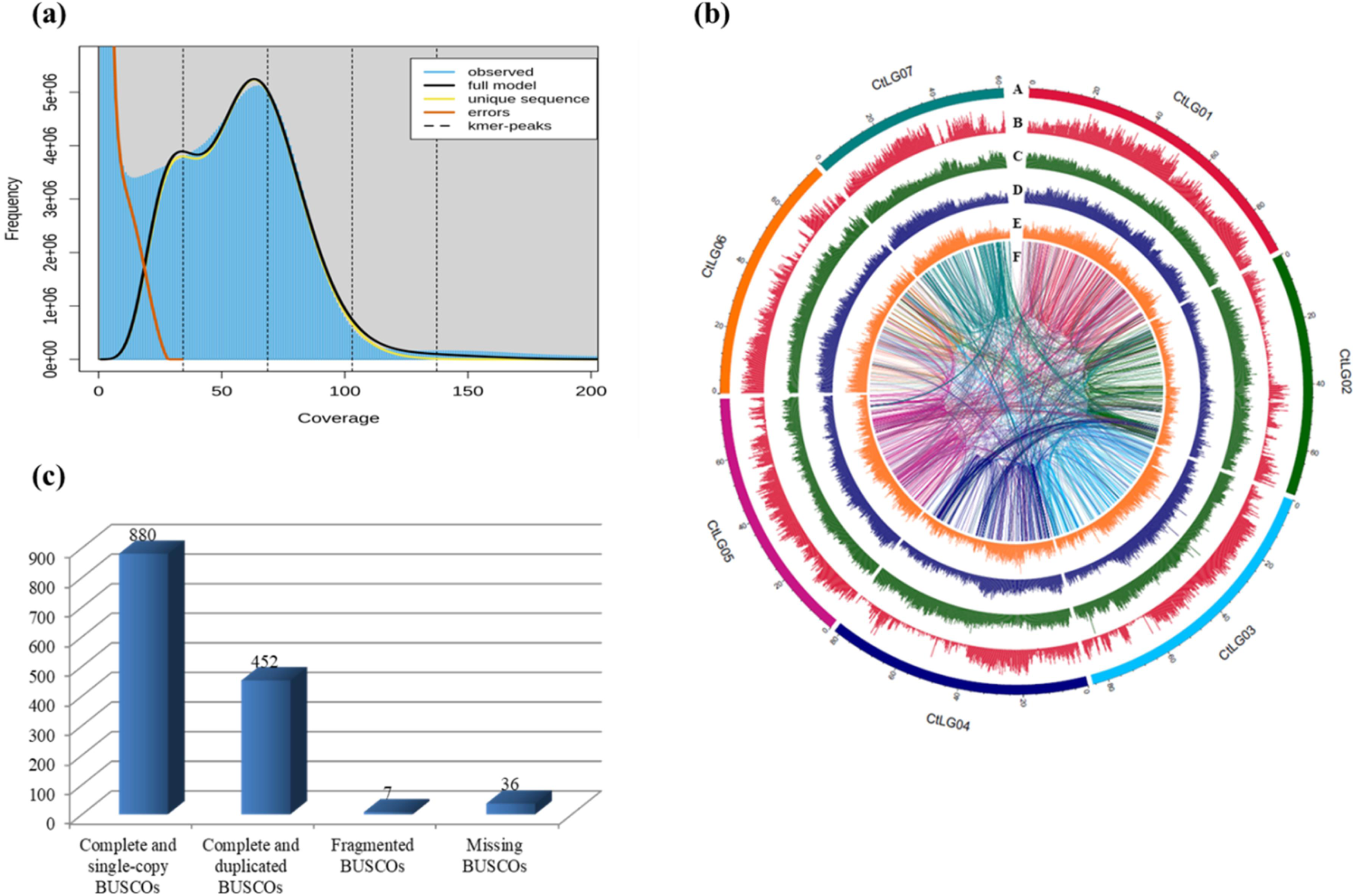
**(a)** The 31-mer frequency distributions of the sequencing reads. The sharp peak on left with low depths represents the random sequencing errors. The middle and right peaks indicate the heterozygous and homozygous peaks, the depths of which are 32 and 65, respectively. **(b)** The cluster bean genome features. Track A represents 7 pseudochromosomes. Track B to E represent the distribution of protein-coding genes, Retrotransposons, DNA elements and Simple sequence repeats, respectively. Track F represents gene duplications in the cluster bean genome. **(c)** Benchmarking Universal Single-Copy Orthologs (BUSCO) analysis of cluster bean annotated genes.

### *De novo* genome assembly

We followed a hybrid assembly approach for developing high-quality chromosome-level genome of cluster bean. Initially, we generated two separate *de novo* assemblies by using 10X genomics linked reads and Oxford Nanopore long reads. We then merged both the assemblies to get minimal sequencing errors and a high degree of sequence continuity for scaffolds.

The raw 10X genomics sequencing data was assembled using Supernova (v2.1.1)^21^, which generates raw assembly as well as scaffold assembly. Removal of vector and mitochondrial contamination was done using seqclean tool through univec vector database (https://www.ncbi.nlm.nih.gov/tools/vecscreen/univec/). We generated a scaffold assembly of 617.76 Mbp comprising 1616 contigs with an N50 size of 4.27Mbp, which was 13.7% longer than the estimated cluster bean genome size.

The long Nanopore sequencing reads were utilized to generate a *de novo* assembly using Canu (v1.6)^22^, which generated 3904 contigs spanning 419.66Mbp with N50 of 775kbp. This primary assembly was further polished with Illumina shotgun and Mate-Pair data to generate an improved assembly of length 441.85Mbp, which comprised of 1548 contigs with N50 of 544.474kbp.

The primary assemblies from 10X supernova and Oxford Nanopore (Canu) were merged with npScarf^23^ and generated a highly contiguous assembly of 550.16Mb comprising 1580 scaffolds with an N50 value of 7.12Mb and longest scaffold of 35.03Mb.

Further, GBS reads of 142 members of a F_2_ population developed from RGC-936/ CAZRI-15-3-8 were aligned against the genome assembly using bwa mapping software and SNP calling was done using Unified Genotyper from the Genome Analysis Toolkit GATK (v3.6). According to UGbs-Flex pipeline, SNPs with allele frequencies <0.1 and >0.9 and adjacent SNPs were discarded. Further markers showing segregation distortion from the expected 1:2:1 Mendelian ratio were discarded on the basis of χ2 test (p < 0.05). We obtained 6113 markers that were imported in JoinMap (v4)^24^, program in Kyazma software package for creating the linkage map containing seven linkage groups (https://www.kyazma.nl/index.php/). Also, the linkage groups were determined at logarithm of odds (LOD) score of 6.0. A total of 1529 scaffolds having 6113 markers in seven linkage groups were merged into specific pseudomolecules using in-house Perl script. Finally, seven genetically anchored pseudomolecules/ chromosomes along with 51 unanchored contigs resulting in the final assembly of 550.30Mb for the clusterbean genome. The chromosome length of the clusterbean genome ranged from 61.32 Mbp (Chr7) to 93.95 Mbp (Chr1) with a scaffold N50 of 78.27Mb **(Table 2)**. The organization of clusterbean genome assembly, gene density, DNA repeat elements, SSRs and duplicate genes were shown in **Figure 1b**. Further analysis is being carried out to improve the assembly to final completion.

**Table 2:** Genome assembly and gene annotation statistics of the cluster bean genome.

| **Assembly features** |  |
| --- | --- |
| Number of scaffolds | 58 |
| Total length of the assembly (bp) | 550,317,005 |
| GC content (%) | 32.64 |
| Scaffold (pseudomolecules) N50 | 78,271,499 |
| Longest scaffold (pseudomolecule) | 93,956,920 |
| Number of contigs | 1580 |
| Max contig length | 35,034,496 |
| Min contig length | 970 |
| Contig N50 | 7,127,297 |
| **Gene models** |  |
| Number of gene models | 34680 |
| Mean transcript length | 3823.77 |
| Mean coding sequence length | 289.115 |
| Mean number of exons per gene | 5.35043 |
| Mean exon length | 249.843 |
| Mean intron length | 681.044 |
| Number of genes annotated | 28,955 |
| Number of genes unannotated | 5725 |

### Identification and annotation of repetitive DNA sequences

For identification and annotation of repetitive DNA sequences in the cluster bean genome assembly, we used a *de novo* repeat library and Dfam^25^ (v3.1) database. First, RepeatModeler (v1.0.10, http://www.repeatmasker.org/RepeatModeler/) was employed to make a *de novo* repeat library of cluster bean genome assembly. Next, we annotated the cluster bean *de novo* repeat library by using Repeatmasker (v4.0.7). Then BLASTn search was performed to annotate unclassified elements from Repeatmasker with the repetitive elements in the Dfam database (https://www.dfam.org/releases/Dfam_3.1/). We identified 582339 DNA repeat sequences covering 42.14% of the cluster bean genome. The most abundant repetitive element type was the retrotransposons making up 29.73% of the genome, including 8.08% of LINES, 21.62% of long terminal repeats (LTRs), 23.3% of DNA transposons and 5.56% of unclassified repetitive elements **(Table 3)**.

**Table 3:**
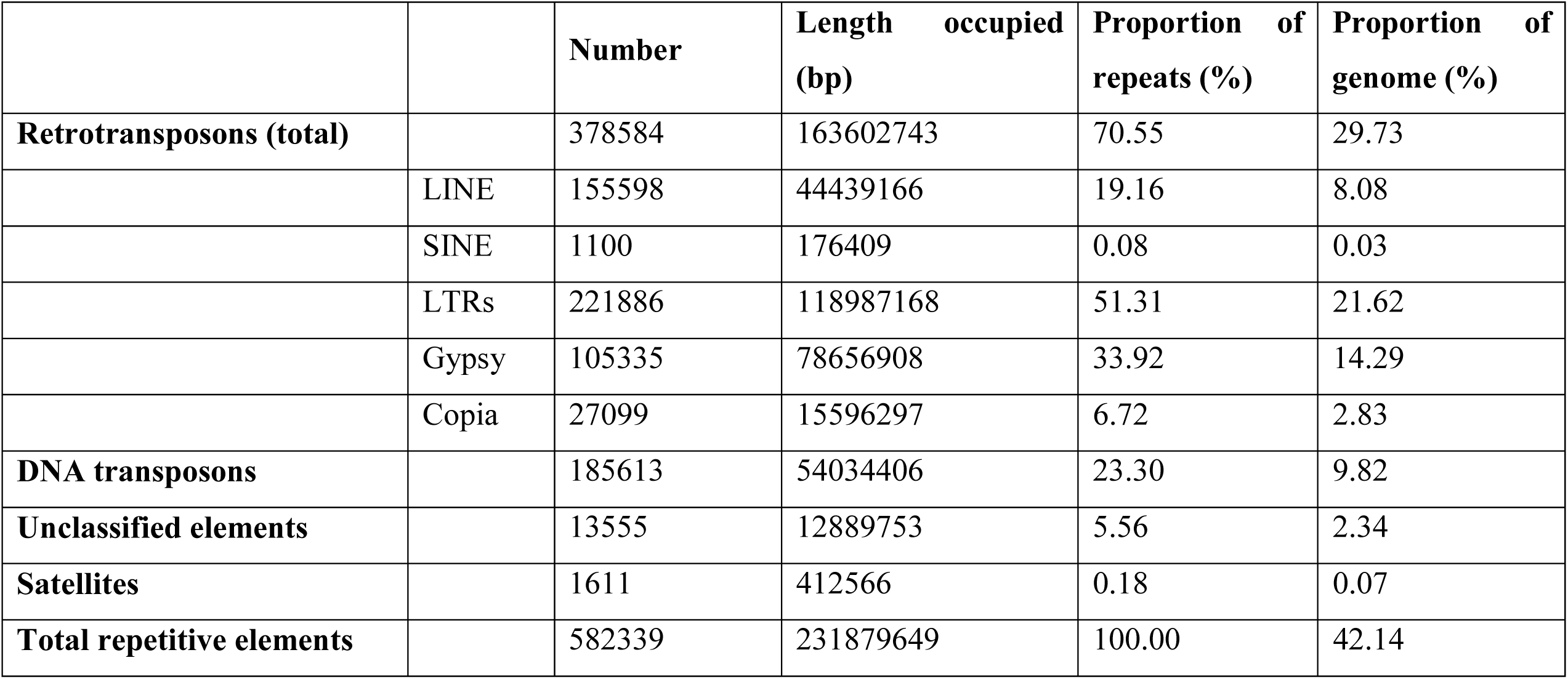
Organization of repetitive elements in the cluster bean genome.

Simple sequence repeats (SSRs) or microsatellites in the cluster bean genome were identified by MISA^26^ program with the following parameters: monomer (n ≥ 10), dimer (n ≥ 6), trimer (n ≥ 5), tetramer (n ≥ 5), pentamer (n ≥ 5), and hexamer (n ≥ 5). A total of 238176 SSRs, covering 0.46% (2.52 Mb) of the cluster bean genome were detected. Among the SSRs, monomers were the most abundant type (71.57%), followed by dimer (16.91%), trimer (9.84%), tetramer (1.31%), pentamer (0.20%), and hexamer (0.16%), respectively.

### Gene model prediction and functional annotation

From the cluster bean genome assembly, the protein-coding genes were predicted using the Seqping (version 0.1.45)^27^ pipeline. Seqping provides species-specific, unbiased gene predictions, thus most suitable for gene prediction of non-model plant genomes like cluster bean. The RNA sequencing data generated earlier by us^15^ was used for transcript assembly with Trinity (version 2.1.1)^28^ and utilized in the Seqping pipeline. Initially, Seqping predicted 37509 protein-coding genes and after clustering with CD-HIT (version 4.6)^29^, 34680 non-redundant gene predictions were obtained.

The efficiency of the gene prediction was evaluated with Benchmarking Universal Single-Copy Orthologs (BUSCO) analysis, using the Plant embryophyta_odb10 lineage with Arabidopsis species (BUSCO v3.1.0)^30^. The BUSCO reported 96.90% of gene predictions as complete including 64% complete single-copy and 32.9% duplicated genes, 2.6% of missing gene models and 0.5% of fragmented gene models **(Figure 1c)**, indicating a high efficiency of gene prediction.

The functional annotation and Gene ontology (GO) terms for each predicted gene model were allocated via InterProScan 5 (version 5.25-64.0)^31^ and Blast2GO (version 4.1)^32^ respectively. About 28955 (78.93%) genes were annotated successfully.

### Identification and annotation of Non-coding RNA genes

Non-coding RNAs (ncRNAs) including tRNAs, rRNAs, and snRNAs in cluster bean genome were identified and annotated using various software packages and databases. First, tRNAscan-SE (version 1.3.1) with default parameters was used to identify and annotate tRNAs and their secondary structures. Total 474 tRNA genes corresponding to a total length of 33.4 kbp were identified. Further, to annotate the ribosomal RNA (rRNA) and small nuclear RNA (snRNA), BLASTn search against the Rfam database (version 14.1) was performed. We found 922 rRNA genes with a total length of 654.07 kbp and 347 snRNA, with a total length of 34.8 kbp.

### Comparative Genome analysis

We used OrthoMCL^33^ (https://orthomcl.org/) to identify ortholog genes among cluster bean and other important crop and model plants including, *Glycine max, Cajanus cajan, Cicer arietinum, Arabidopsis thaliana, and* a monocot *Oryza sativa*. The details of orthologous gene families are mentioned in **Table 4**.

**Table 4:** Gene family analysis of the predicted cluster bean genes in comparison to other plant genomes.

| **Species** | **No. of predicted genes** | **No. of Genes in Orthologous groups** | **No. of Genes not in orthologous groups** | **Total no. of orthologous groups** | **No. of Inparalogs genes** | **No. of Single copy Orthologs** | **Average genes group** |
| --- | --- | --- | --- | --- | --- | --- | --- |
| *C. tetragonoloba* | **34680** | **13584** | **21096** | **10639** | **3356** | **8555** | **1.28** |
| *G. max* | 56044 | 43158 | 12886 | 24376 | 34900 | 5555 | 1.77 |
| *C. cajan* | 31841 | 23355 | 8486 | 20812 | 9329 | 12521 | 1.12 |
| *C. arietinum* | 26164 | 15505 | 10659 | 11639 | 4835 | 9939 | 1.33 |
| *O. sativa* | 55986 | 25591 | 30395 | 14108 | 19797 | 5522 | 1.81 |
| *A. thaliana* | 28775 | 21830 | 6945 | 14447 | 14471 | 6908 | 1.51 |

### Data Records

The Guar *(Cyamopsis tetragonoloba)* genome assembly data is deposited in NCBI under BioProject and will be made available soon.

### Code availability

The bioinformatics tools/packages used in this research are described below along with their versions, settings and parameters.

**1) Supernova:** version 2.1.1, default parameters; **(2) Canu:** version 1.6, default parameters; **(3) Seqclean:** with default input sequence in fasta format -v /usr/db/adapters, /usr/db/UniVec, /usr/db/linkers\,-s/usr/db/ecoli_genome,/usr/db/mito_ribo_seqs; **(4) npScarf(jsa**.**np**.**npscarf):** default parameters; **(5) UGbs-Flex pipeline:** with available python and perl script; **(6) GATK**: version 3.6; **(7) BUSCO:** version 3.1.0, -l eukaryota_odb10, -e 1e-05, --augustus_species Arabidopsis; **(8) RepeatModeler:** version 1.0.10; **(9) RepeatMasker:** version 4.0.7, search engine= rmblast; **(10) Jellyfish:** version 2.0; **(11) GenomeScope**: version 2.0; **(12) MISA perl script**:; **(13) Rfam**: version14.1, (January 2019, 3016 families); **(14) Seqping**: version 1.45, -e 1e-10; **(15) GlimmerHMM**: version 3.0.4; **(16) Maker:** version 2.31.9; **(17) tRNAscan-SE**: version 1.3.1; **(18) CD-HIT:** version 4.6, default parameters; **(19) Trinity:** version 2.1.1, -- seqType fq --full_cleanup --min_contig_length 250; **(20) InterProScan 5:** version 5.25-64.0; **(21) Dfam:** version 3.1 (June 2019, 6915 entries); **(22) JoinMap:** version 4, default parameters; **(23) OrthoMCL:** version 1.2

## Acknowledgements

Clusterbean Sequencing Initiative was funded completely by ICAR-CRP on Genomics. We are grateful to Dr J. K. Jena (Coordinator & DDG (FS), ICAR, New Delhi, India and Dr Vindhya Mohindra (Co-Coordinator, CRPG, NBFGR, Lucknow, India) for providing constant support, encouragement and efficient coordination of the entire program objectives. The authors acknowledge Dr T Mohapatra (Secretary, DARE & DG, ICAR, New Delhi) for constantly encouraging and giving critical suggestions for improvement of the program activities.

